# How Homeostasis Limits Keratinocyte Evolution

**DOI:** 10.1101/548131

**Authors:** Ryan O. Schenck, Eunjung Kim, Rafael R. Bravo, Jeffrey West, Simon Leedham, Darryl Shibata, Alexander R.A. Anderson

**Affiliations:** Integrated Mathematical Oncology Department, H. Lee Moffitt Cancer Center and Research Institute, Tampa, FL 33612, USA; Wellcome Centre for Human Genetics, University of Oxford, Oxford, OX37BN, UK; Department of Pathology, Keck School of Medicine of the University of Southern California, Los Angeles, California 90033, USA

## Abstract

The skin is the largest human organ, functioning to serve as the protective barrier to the harsh, outside world. Recent studies have revealed that large numbers of somatic mutations accumulate, which can be used to infer normal human skin cell dynamics^1-6^. Here we present the first realistic mechanistic epidermal model, that uses the ‘Gattaca’ method to incorporate cell-genomes, that shows homeostasis imposes a characteristic log-linear subclone size distribution for both neutral and driver mutations, where the largest skin subclones are the oldest subclones. Because homeostasis inherently limits proliferation and therefore clonal sweeps, selection for driver mutations (NOTCH1 and TP53) in normal epidermis is instead conferred by greater persistence, which leads to larger subclone sizes. These results reveal how driver mutations may persist and expand in normal epidermis while highlighting how the integration of mechanistic modeling with genomic data provides novel insights into evolutionary cell dynamics of normal human homeostatic tissues.

---

Recent studies have documented that substantial numbers of mutations accumulate in normal human tissues, with evidence for selection of canonical, oncogenic driver mutations^2-7^. Typically, selection for these driver mutations is modeled by increased cell proliferation; however, requirements of skin homeostasis imposes spatial constraints that inherently limit selective sweeps. Because these mutations do not appear to disrupt homeostasis, these mutations can also provide important new information on normal human tissue dynamics, which are typically studied with experimental fate markers in animal systems.

Although substantial numbers of mutations accumulate with age, and a quarter of all cells carry driver mutations^2^, skin thickness and mitotic activity do not significantly change. A controversy currently exists between our understanding of mutation selection and neutral evolution within tissues^1,2,8,9^, in part because our understanding of how a mutant clone is able to expand within normal tissue is completely lacking. Furthermore, most mutations are passengers and skin subclone size/frequency distributions are consistent with neutral evolution or the absence of detectable selection. There is strong evidence of selection by driver mutations (NOTCH and TP53) manifested by dN/dS ratios and their generally larger subclone sizes in normal skin^2^. Varying exposures to sunlight between individuals also complicates our understanding of mutation accumulation. The challenge is to develop a model that integrates selection, neutral evolution, and sunlight. The reward would be a realistic model of lifelong human skin cell dynamics.

In the scope of the epidermis, first principles dictate a constant cell number through equal loss and replacement of cells, a constant tissue height, and constant stem cell numbers. Using these principles, we built a three-dimensional mechanistic model of the human epidermis using HAL^10^, a comprehensive cell based modeling framework allowing users to focus on the development of model specific functionalities. We use this model to infer how both neutral and driver mutations accumulate and exhibit clonal expansions without disrupting homeostasis (Figure 1). We based our model on a simplified version of the vSkin model^11^ which is capable of wound repair, and uses EGF as a key diffusible stromal growth factor to define a stochastic stem niche within the basal layer and determine keratinocyte birth/death (Figure 1 and Figure S1). In response to this EGF we parameterize the model to a progenitor loss/replacement rate of 0.51 week^-1 1^.

**Figure 1.**
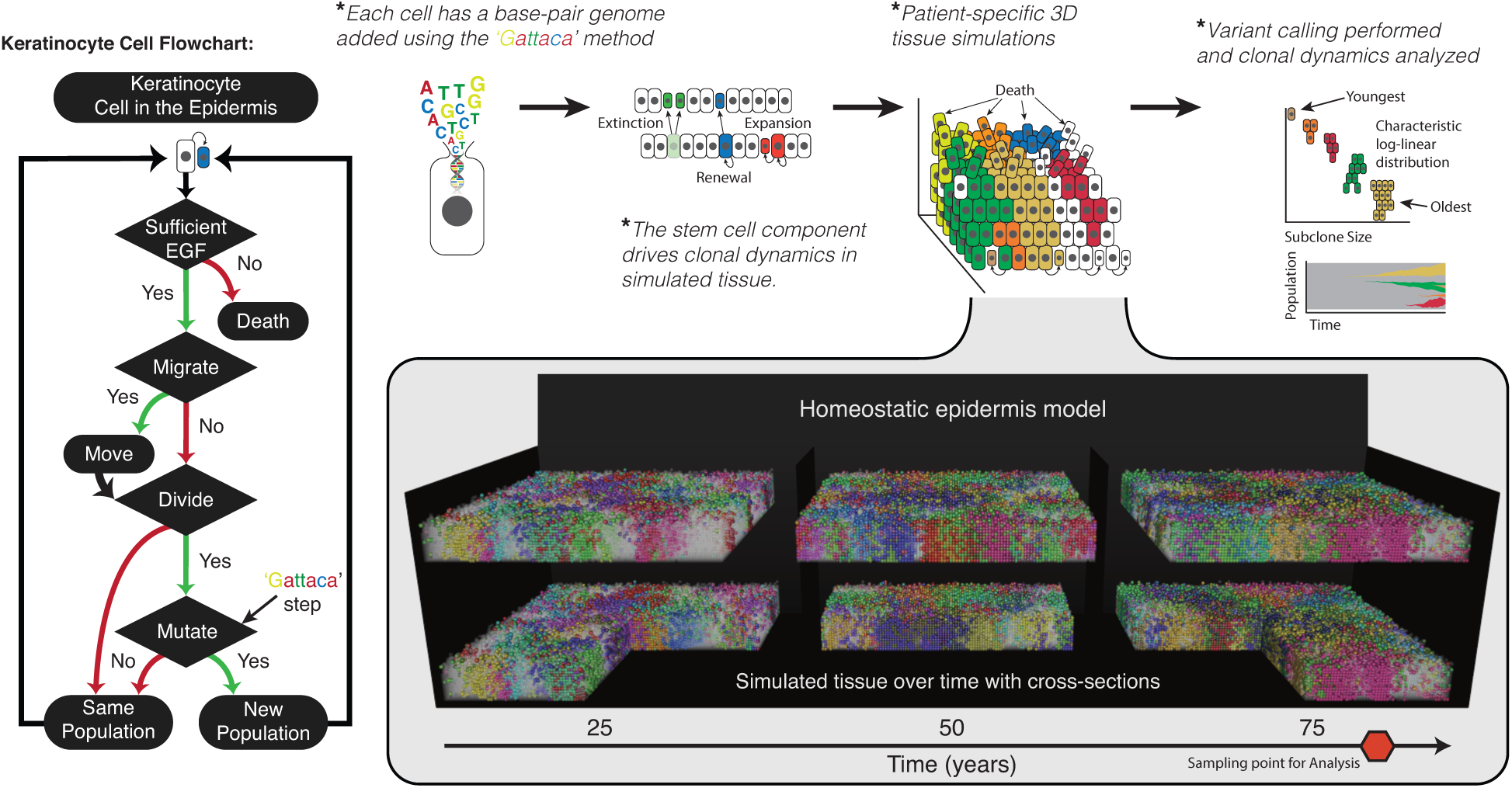
Homeostatic epidermis model with high resolution genomes. The homeostatic epidermis is comprised of individual keratinocyte cells that are governed by a set of decisions (Flowchart). Spatial structure of the three-dimensional (3D) model and an EGF gradient governs the loss/replacement in the stem cell niche at the basal layer. The model is entangled with DNA sequence data by the Gattaca step, where base-pair mutations can be introduced with every *in-silico* cell division. These mutations act as cell fate markers and different sub clones are represented by different cell colors. Therefore, cells in the simulated skin produce a patchwork of different sized sub clones, where each colored population differs by at least one mutation.

In a model where stem behavior is a niche driven phenomenon, the target loss/replacement rate in a 1mm^2^ area with one in three cells being self-renewing becomes 0.1683 week^-1^ overall (basal cell density of 10,000 cells). Meaning that any cell within the basal layer possesses an equal probability of assuming a self-renewing role during a given time step. Within this system transient cell divisions propagate superficially in response to the remaining EGF diffusible. However, these transient divisions contribute minimally to the overall loss/replacement in the system, where 39.8 ± 0.85% occurs within the basal layer and diminishes rapidly within each layer approaching the *stratum cornea* (Figure S1B). Overall population size (mean cell number of ≈ 120,000 within a simulated 1mm^2^ biopsy) is balanced by death where the top layers of the epidermis transition into the *stratum cornea* due to low EGF concentrations.

To assess clonal expansions and mutational data for comparison with patient data we developed the ‘Gattaca method’ (see Methods for full details), where 72 specific genes, tracked at base pair resolution, with gene specific mutation rates are imbedded within each keratinocyte, accurately capturing the mutation distribution consistent with UV exposure (Figure 1B). Gattaca stands for the four DNA bases, GATC, and reflects that individual base changes can be used as intrinsic cell fate markers that can be interrogated by DNA sequencing of human cells. This method, used within each cell, allows us to fully represent the tissue architecture of a homeostatic epidermis with realistic mutation accumulation (Figure 1 and Supplemental Movie S1).

Simulations of variable biopsy sizes (0.75, 1.0, and 0.1*θmm*^2^), age matched to comparable patients, reconstructs a log-linear subclone size-age distribution with neutral mutations that easily captures the extent of clone size variability within patients (Figures 2A and 2B). We see that small clones dominate the highest frequencies while a heavy right tailed distribution is reflected by the clonal exponential size dependency (Figures 2A and 2B) consistent with patient data^1,3,12^. We then perform pairwise Kolmogrov Smirnov tests on each patient and the simulated biopsy’s First Incomplete Moment distribution (FIM). We fail to reject the null hypothesis, that the distribution of clone sizes are from different distributions, for the vast majority of our simulation versus patient biopsy comparisons, but are able to reject the null for a select few comparisons (Figure 2B). However, for all patient biopsies at least one of the patient specific simulations fails to reject the null hypothesis. In a homeostatic epidermis, most subclones rife with non-functional (i.e. neutral) mutations are small, recent, and destined for extinction due to random stem cell niche turnover (Figures 2A, 2B, and 3E). Persistent, older subclones are rarer and larger (Figure 3D). Hence, the observed log-linear subclone mutation size distributions (Figure 2B) are an emergent property of the homeostatic constraints of normal human skin. These results reveal that it is highly improbable that a mutation is capable of expanding larger than any other, given homeostasis. Age-related clonal size dependencies exist even for advantageous driver mutations in the homeostatic system (Figures 3D-E). This implicates the constraints of the homeostatic tissue architecture as the primary factor in defining the age-related exponential size dependency.

**Figure 2.**
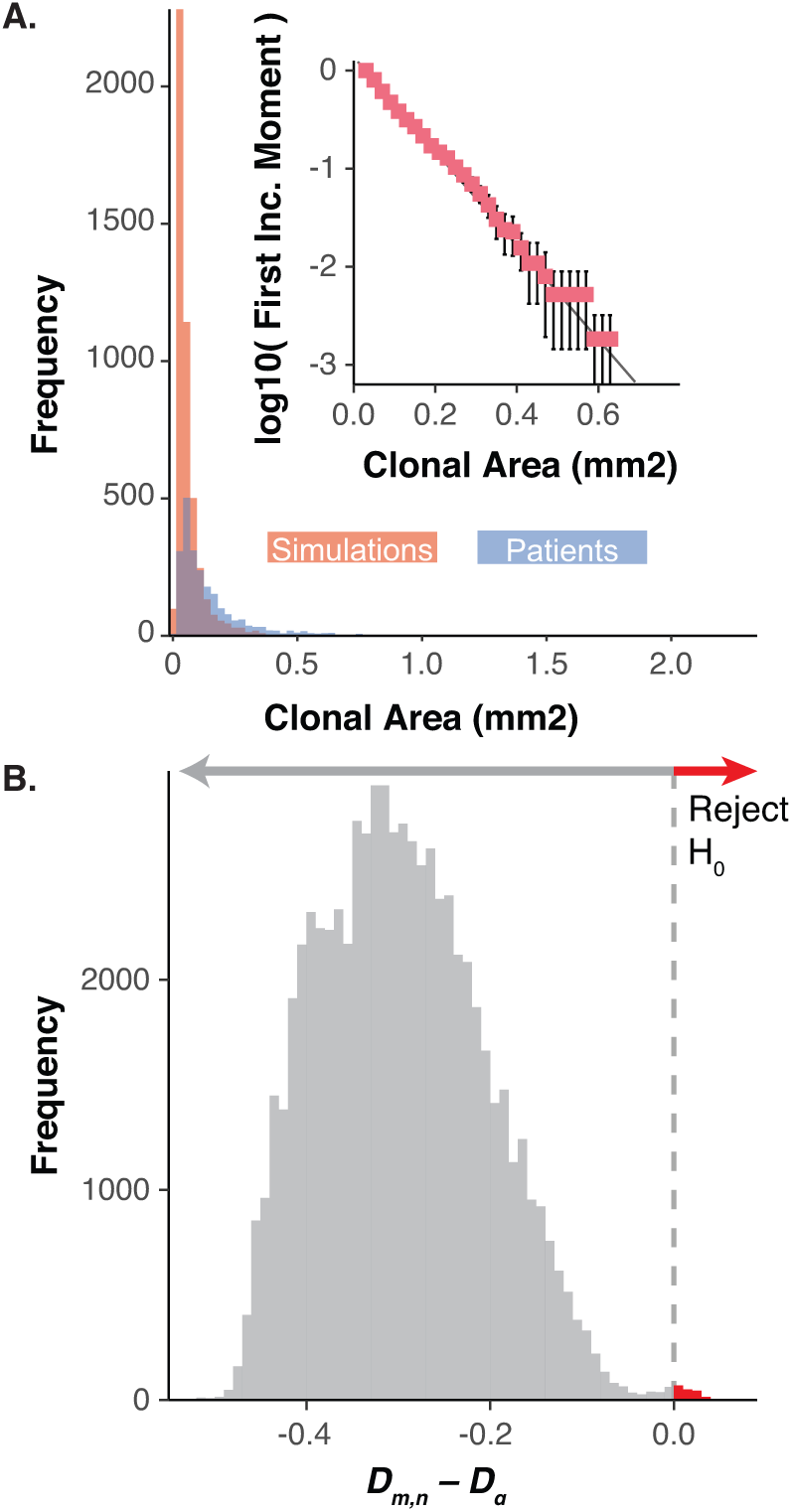
Homeostasis imposes log-linear clonal distributions. Neutral model dynamics from three-dimensional (3D) simulations of various sizes where (A) is the cumulative clonal area frequencies of all patient biopsies (blue) and randomly chosen simulations for each of the patient specific biopsies such that the number of simulations chosen is equal across the three simulated sizes and the sum of the number of biopsies equals that of the total biopsies for the patient (red). The inset figure shows the log-10 transformed first incomplete moment for the same random sampling of patient comparable simulations for PD20399 (error bars denote standard deviation for 100 repeated samplings). (B) Shows the difference from the Komlogrov-Smirnov test statistic (*D*_*m,n*_) and critical value (*D*_*α*_) for all patient biopsy to patient specific model simulation’s first incomplete moment distributions for all simulation sizes. The red arrow denotes comparisons where the null hypothesis can be rejected.

**Figure 3.**
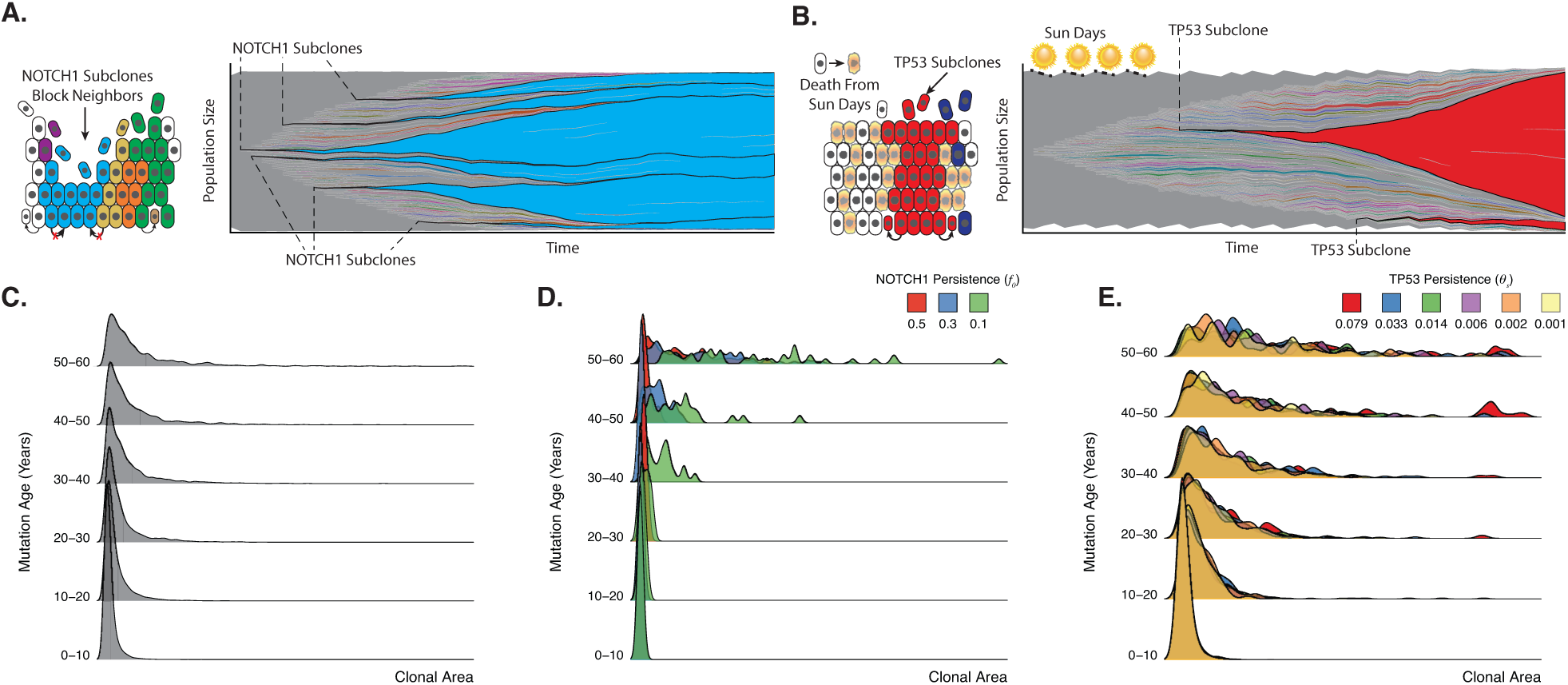
Homeostasis, *a priori*, constrains clones in a functionally heterogeneous tissue, dictating an age-dependent clonal expansion. Clonal dynamics in homeostasis are a function of complex interactions between the microenvironment and the external environment. During homeostasis, every cell is equal relative to its neighbors, i.e. neutral. NOTCH1 disrupts neutral dynamics by offering a slight advantage via a blocking probability (*f*_0_) which prevents neighboring cells from dividing into their positions (A schematic). Whereas, TP53 mutations aren’t subject to UV damage, when a proportion (*θ*_*s*_) of non-TP53 mutant cells may be killed by UV damage (B schematic). (A) and (B) next to each schematic, show clonal expansions for a single simulation for NOTCH1 advantaged clones, blue, and TP53 advantaged clones, red, in a 1mm^2^ simulation to 58 years, respectively. The age dependence is shown in the density plots broken by mutation age for 100 replicate simulations up to 58 years for neutral (C), NOTCH1, with varying strengths (*f*_0_) (D), and TP53, with varying strength (*θ*_*s*_) (E).

Previously, NOTCH1 and TP53 mutations were observed to have larger subclone sizes relative to neutral mutations^2^, indicating positive selection. To better define this selection within the context of homeostasis, we simulated increases in the ability of NOTCH1 mutant subclones to displace neighboring cells, which leads to clonal distributions consistent with positive selection (Figure 3E). However, the requirement to maintain homeostasis indicates this mechanism comes with a cost (Figures 3 and 4). The stronger the advantage, the greater the loss of homeostasis, creating a catch-22 where a strong NOTCH1 mutant clone with too much of a selective advantage destroys the epidermis. Therefore, young, rapidly expanding large NOTCH1 subclones are never observed (Figures 3 and 5). This is consistent with the observation that patient NOTCH1 mutations are smaller but more frequent than observed TP53 mutations^2,3^. Instead, similar to the neutral mutations, the largest NOTCH1 subclones are the oldest NOTCH1 mutations. Although driver mutations can only confer small selective advantages without disrupting homeostasis, even small persistence advantages expressed at earlier patient ages pays dividends and inherently leads to larger and more frequent NOTCH1 subclones in adult skin (relative to neutral mutations).

**Figure 4.**
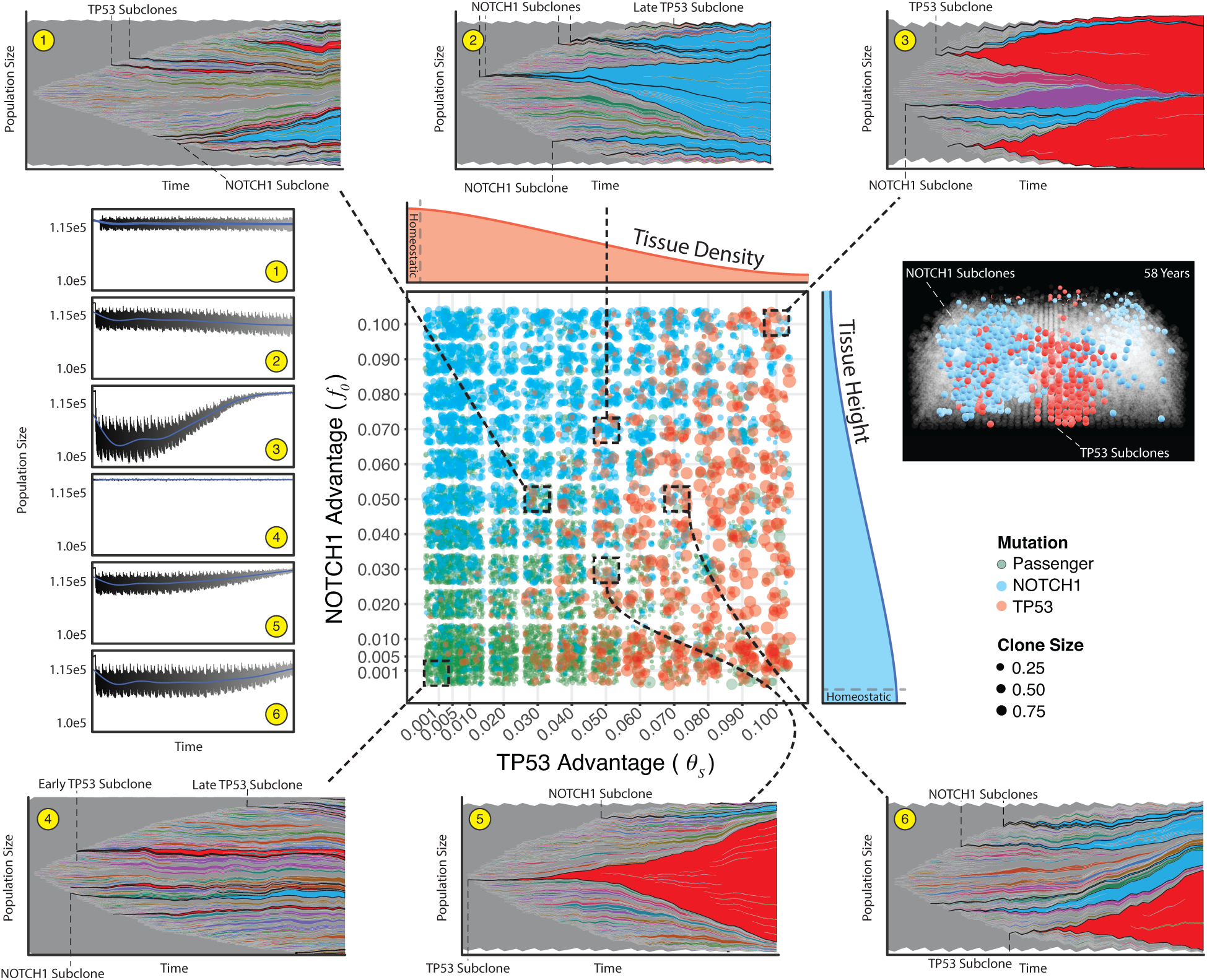
Combined effects reveal homeostatic recovery and recapitulate patient clone distributions. The primary, central, plot reveals clone sizes for 1*mm*^2^ simulations for four different ages with five replicates across a manually defined scale of parameter pairs for NOTCH1 (*f*_0_) and TP53 (*Θ*_*s*_) advantages (n=2,880 simulations). Density subplots show an idealized example of how each mutation type effects homeostasis (red and blue for TP53 and NOTCH1, respectively). The dashed boxes and lines show the evolutionary dynamics for a single simulation for a 58 year simulation (population frequencies greater than 0.001) for six parameter pairs and the corresponding overall population sizes over time (1-6 yellow circles, blue line is a locally weighted smoothing line). For a single parameter pair (*Θ*_*s*_ = 0.003, *f*_0_ = 0.05) a 0.4*mm*^2^ tissue is shown where only NOTCH1 (blue) and TP53 (red) clones are colored.

**Figure 5.**
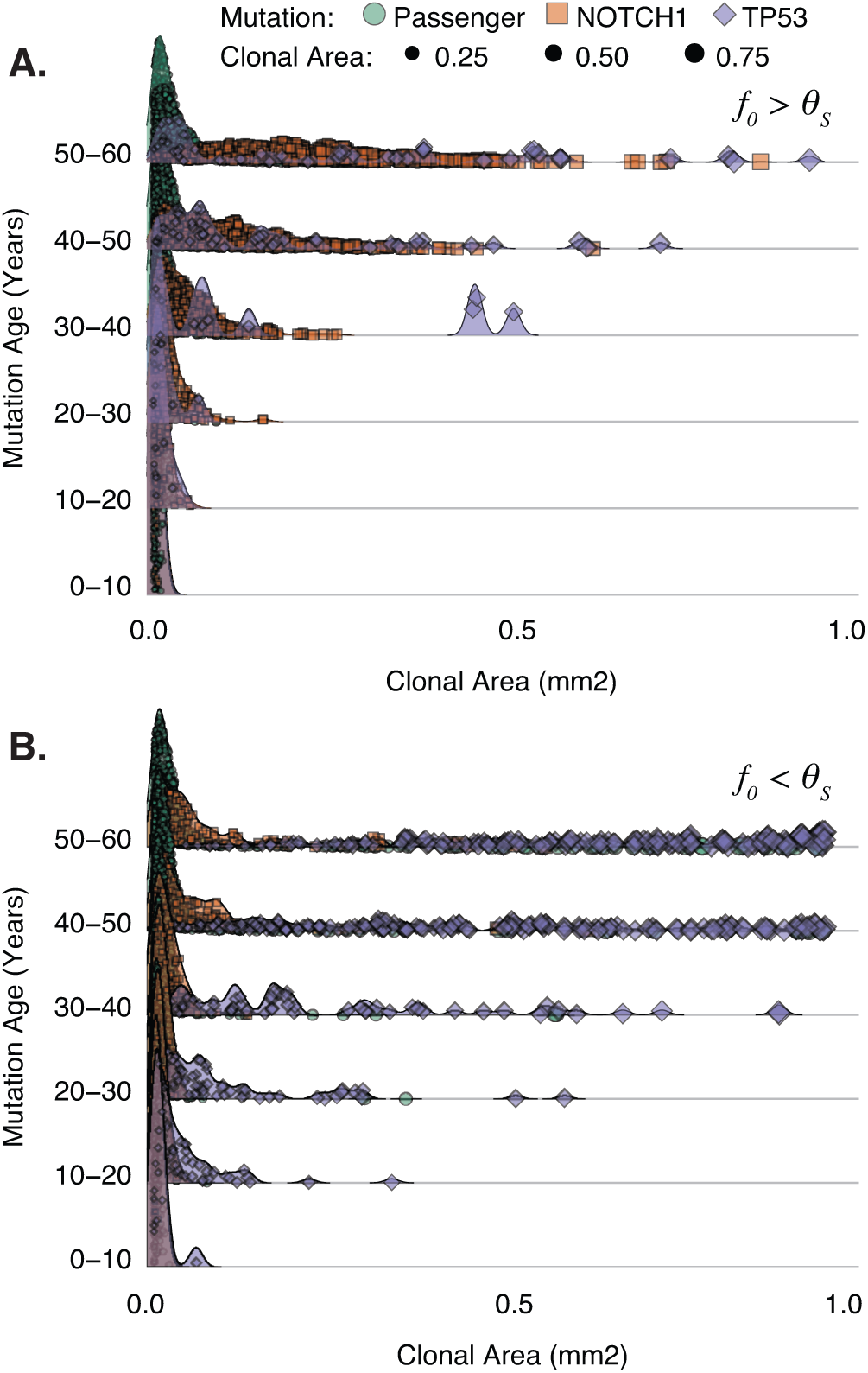
Combined effects adhere to age-dependent expansions. All 58 year 1*mm*^2^ paired parameter simulations from Figure 3 are displayed where TP53 persistence (*Θ*_*s*_) is greater than NOTCH1 persistence (*f*_0_) (A) and *f*_0 > *Θ*_*s*_ (B). Individual clones are broken into three groups (passenger, NOTCH1, and TP53) with corresponding sizes of each clone.

TP53 mutations protect against cell death and require sunlight for selection because TP53 subclones are smaller in non-sun exposed skin^13^. Normal sun-exposed skin does not exist because the extent of sun exposure varies massively between patients. Here we incorporate this important environmental variability through “sun days”, where our homeostatic tissue takes a sun-filled vacation and a proportion of non-TP53 mutant cells are killed by UV irradiation (Figure 3B). This mechanism, unlike NOTCH1, renders young TP53 capable of rapid expansion as the skin is exposed to more sun days (Figure 3E). However, similar to NOTCH1, simulated TP53 subclonal expansions are also constrained by homeostasis, which is disrupted by too much UV exposure leading to a loss in tissue density.

When we shed light on these functional differences of NOTCH1 and TP53 existing within the same simulated biopsy, the observed patient data is further understood (Figures 4 and 5). The key variable in tissue homeostasis and clonal expansions is sun exposure. In the absence of sunlight, TP53 mutations are subject to drift, along with all other mutations. Consistent with patient data we see more frequent occurrences of NOTCH1 mutations at smaller sizes across a range of fitness values. In this same tissue TP53 mutations occur less often but are able to expand to a larger area upon sun exposure, even when they arise later, thereby shifting the age-dependent distribution to a more recent point in time. In rare occurrences we see passenger mutations carried to larger clonal areas, but they are largely relegated to the lowest frequencies (Figure 4). As with NOTCH1 and TP53 in isolation, we see that in a tissue where both exist, homeostasis constrains these clones from expanding. However, entirely unexpected, we see that with the presence of a NOTCH1 mutation(s) and TP53 mutation(s), sunlight can minimize the effects of a NOTCH1 mutations (i.e. reduce subclone size) while returning the tissue to a homeostatic state. This is because as a TP53 mutation expands with UV irradiation, the population size recovers over time (Figure 4). Even with the presence of sunlight and both mutations we observe an age-dependent log-linear relationship where only the earliest mutations expand to an appreciable size within the tissue (Figure 5).

Here we have presented the first data-driven *in silico* model able to capture the homeostatic cell dynamics of the human epidermis with high-resolution base pair tracking using the ‘Gattaca’ method embedded within the HAL^10^ framework. Our *in silico* epidermal model reveals that mutational persistence is the key to observed subclone frequency-size distributions. Strikingly, both neutral and driver mutations exhibit similar frequency-size distributions consistent with neutral evolution, reflecting that random stem cell turnover is a feature of normal epithelial homeostasis. Although mitotic stem cells accumulate mutations, random stem cell death with replacement leads to mutation flushing such that most mutations are lost or are limited to small subclones. This work broadens our current understanding of selection and fitness acting in a homeostatic, normal tissue, where selection reflects persistence rather than selective sweeps, leading to larger subclone sizes in adult skin. The next time you take that selfie of your homeostatic epidermis on a tropical, sun filled vacation, think about the portrait of the mutations you would likely observe and the mutational landscape present just at the surface.

## Acknowledgements

ROS is supported by the Wellcome Trust (grant no. 108861/7/15/7) and the Wellcome Centre for Human Genetics (grant no. 203141/7/16/7). JW, EK and ARAA are supported by Physical Sciences Oncology Network (PSON) grant from the National Cancer Institute (grant no. U54CA193489). SL is supported by the Wellcome Trust (grant no. 206314/Z/17/Z). DS is supported by the Cancer Systems Biology Consortium grant from the National Cancer Institute (grant no. U54CA217376 and grant no. P01 CA196569). ARAA is also supported the Cancer Systems Biology Consortium grant from the National Cancer Institute (grant no. U01CA23238) and the Moffitt Cancer Center of Excellence for Evolutionary Therapy.

## Author Contributions

ROS, DS and ARAA conceived the question and model design. ROS wrote the model, performed analysis of model outputs, and wrote the manuscript with critical comments and inputs from SL, ARAA, and DS. EK and RRB assisted with editing the manuscript and model design. JW assisted with manuscript edits and select data visualizations. All authors have read and edited the final manuscript.

## Methods

Here we have constructed a homeostatic, three-dimensional, hybrid cellular automaton (3D-HCA) to mechanistically model the human epidermis using the HAL framework^10^. We based our model on a simplified version of the vSkin model we previously developed^11^ to investigate the clonal dynamics of keratinocyte populations (Figure S1). In this simpler model, while we model keratinocyte dynamics, we do not explicitly consider melanocytes and stromal cells (not in the epidermis) but rather, a single diffusible growth factor acting to define the stem niche on the basal layer. We chose EGF, as a key growth factor, to determine keratinocyte birth/death (Figure S1 and Supplementary Information). We define our area to approximate different sized biopsies assuming a median cell size of approximately 15µ*m*.

### Microenvironmental factor

The microenvironmental factor, epidermis growth factor (*EGF*), governing keratinocyte apoptosis and proliferation is defined by a partial differential equation that dictates how *EGF* diffuses throughout the system over time. Similar to Kim *et al*.^11^ & Basanta *et al*.^14^, due to the interactions between discrete, on lattice cells, and the continuous *EGF* variable we describe the discretized version of the equation. Over time EGF diffuses from an imposed, unchanging source (*E*_*C*_) along the basal layer (i.e. *E*(*x, y*, 0) = *E*_*C*_) with a diffusion rate (*ψ*, Table S1), representative of fibroblast production from the dermis, is consumed by keratinocytes (*K*) at a consumption rate (*γ*, Table S1), and decays naturally (*λ*, Table S1). The temporal from time *t* to time *t* + *δt* of *EGF*([*E*(*η, t*)]), in its discretized form using a first order Euler method, on a three-dimensional lattice is defined as follows:

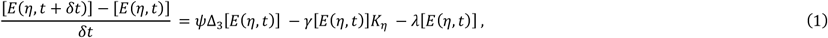

Where *η* ≡ (*η*_*x*_, *η*_*y*_, *η*_*z*_), *δ*_*t*_ denotes the time step, *K*_*η*_ is set to 1 if the lattice point *η* is occupied by a keratinocyte and 0 otherwise. The Δ_3_ denotes the central difference approximation of the Laplacian operator in a three-dimensional space:

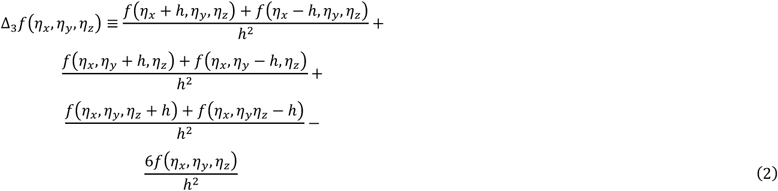

where *h* is the grid size. Boundary conditions on the top (*η*_*z*_ = *Z, Z*: height) are no-flux 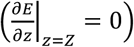 and on the bottom (*η*_*z*_ = 0) are Dirichlet (*E*(*η*_*x*_, *η*_*y*_, 0) = *E*_*C*_), while periodic boundary conditions are present on the left & right and front & back (*E*(0, *η*_*y*_, *η*_*z*_)= *E*(*X, η*_*y*_, *η*_*z*_) & *E*(*η*_*x*_, 0, *η*_*z*_)= *E*(*η*_*x*_, *Y, η*_*z*_)). Each time step we solve equation (1) until steady state reached (<100 iterations). A steady state value of EGF is used to determine keratinocyte birth and death in the model.

### Keratinocyte dynamics model

Cell death can occur one of two ways at each time step for each cell (cell time step is equal to 1 day). First, the cell is dependent upon EGF concentration to prevent a cell from undergoing apoptosis. This value, the level at which apoptosis can occur by chance (*α*, Table S1), is a model specific parameter and translates to a cell undergoing apoptosis when a randomly selected number *C*_*α*_ ∈ *U*(0,1), U: uniform distribution (0,1), is less than 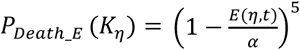, (i.e., if 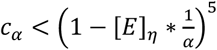, the keratinocyte *K*_*η*_ at a lattice point *η* dies).

If a cell survives in low EGF concentrations or is not in an area with sufficiently low EGF it is then subject to random death representative of an intrinsic probability of death (*P*_*Death_i*_(*K*_*η*_) = *θ*) where death will occur if a randomly selected number from (0,1), *c*_*θ*_ ∈ *U*(0,1), is less than *θ* (i.e., if *c*_*θ*_ *< θ, c*_*θ*_ *∈ U*(0,1), then death occurs).

Provided a cell does not die within a given time step it is capable of moving to an empty lattice position in its 6 immediate nearest neighbors i.e. the von Neumann neighborhood (*N*_(*x,y,z*)_^*v*^ = {(*x,y,z* + *h*),(*x,y,z* − *h*),(*x* + *h,y,z*),(*x* − *h,y,z*), (*x,y* − *h,z*),(*x,y* + *h,z*)}). This location is determined by assessing if any empty lattice positions exist within the cell’s neighborhood. If multiple spaces are empty, one is chosen randomly. The model specific parameter dictating the probability of movement (*P*_*Move*_(K_*η*_) = *ρ*, Table S1) governs the cells ability to move into that empty space so that when a randomly selected number *c*_*ρ*_from (0,1) is greater than *ρ*, then the cell moves into that empty position (i.e., if c *c*_*ρ*_(∈ *U*(0,1)) *≥ ρ*, the cell will move into a randomly selected empty neighborhood, *N*_(*x,y,z*)_^*v*^).

Cells divide based on the underlying EGF concentrations (*P*_*birth*_ (*K*_*η*_) = *ξ*[*E*]_*η*_), where *ξ* is the proliferation scale factor (Table S1). This results in a stem–like population of cells dependent on the underlying EGF concentration. A cell (*K*_*η*_ at *η*) may divide into two identical daughter cells (*D*_1,2_) if a randomly selected number *c*_*d*_(*c*_*d*_ ∈ *U*(0,1)) is less than *ξ*[*E*]_*η*_ regardless of there being an empty position. One daughter cell occupies the position of the parent cell (location of *D*_1_ = *η*_*x,y,z*_) while the other daughter cell (*D*_2_)occupies one of five locations, being above or on one of the four orthogonal neighbors of the parent cell. This location is governed by *ω*, the division location probability, whereby the second daughter cell is placed in a position according to

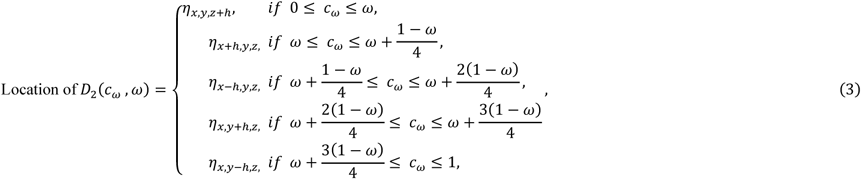

where *η*_*xyz*_ is the coordinates of the parent cell’s lattice position, *h* is the grid size, and *c*_*ω*_ is a randomly selected number from (0,1) (i.e., *c*_*ω*_ ∈ *U*(0,1)). Boundary conditions for cells are the same as those for EGF, being periodic on the left & right and front & back to enable spatial competition, no-flux on the bottom (*η*_*z*_ = 0), and Dirichlet (Cell (*η*_*x*_, *η*_*y*_, *Z* = 0) on the top.

### *Gattaca* Genome Modeling Method

Each cell agent contains a base pair resolution of 72 genes with genomic positional information constructed from the reference hg19 genome using ANNOVAR^15^. Each gene has a gene specific mutation rate such that the mean mutation rate (*μ*_*g*_) is normalized to ≈ 3.2 × 10^−9^ *bp*^−1^ *division*^−1^(likely a conservative estimate, but normal somatic mutation rates can vary wildly^16-19^). Mutations within the rest of the genome are tracked with less resolution. While a random base within any given position can be tracked, these mutations are not of particular interest here, but can be used in statistical analysis and model–data comparisons. However, due to substantial computational requirements to track such large amounts of information for each cell this method is employed sparingly and not in all model simulations. This method allows for a more realistic mutation accrual, since a genome wide mutation rate would be biologically unrealistic due to a large number of covariate factors (e.g. level of expression, location on chromosome, and GC content, etc.) leading to differences in mutation rates^20^. A mutation rate applied uniformly across the genome leads to the longest genes, such as ERBB4, PTPRT and BAI3 accumulating the most mutations (in our model) while NOTCH1-4 would acquire almost no mutations. This is not observed in normal epidermis, where NOTCH1 and a number of various other shorter genes are frequently mutated.

Upon initialization of the model we construct Poisson distributions for each gene given the mutation rate, *μ*_*g*_, and the length of the gene, *L*_g_, providing the expected number of mutations upon division. Where the number of mutations for each gene, *X*_*g*_, is *X*_*g*_ ∼ *poisson* (*μ*_*g*_ × *L*_*g*_),and the specific base affected is determined by a multinomial distribution constructed for each A, C, T, and G allowing us to define and capture a specific mutational signature observed in the data. Keratinocyte (and melanocytes not considered here) contain, disproportionately more C>T transitions^2,3^; this needs to be accounted for by adjusting the probability of a C>T mutation occurring versus all other mutation types. Here we set the probability of a cytosine mutation to 0.5 and split the remaining bases to an equal probability. Finally, we determine the position within the gene by applying a uniform probability to all of the same bases within the gene. All ancestral mutations are inherited by daughter cells. Within the model distributions are constructed using the Colt java library upon initialization.

### Model Parameterization

Parameter values for the spatial model were selected based on iterative redundancy analysis (RDA) using R^21^ (vegan package^22^), where constraining variables were the emergent properties measured as a necessary component of homeostasis (see Supplementary Information):

Primary Constraining Variables:

1. Mean Tissue height
2. Mean Loss/replacement rate within the basal layer

Secondary Constraining Variables:

1. Mean Overall tissue turnover time
2. Mean cell age
3. Mean Basal cell density

During the first iteration of parameterization variables are set randomly choosing values between zero and one for 20,000 two-year simulations. EGF parameters (*ψ,γ,λ*), where values were examined between zero and 1/6, were not subject to this iterative process. For the diffusion coefficient (*ψ*), values greater than 1/6 (for 3D, 0.25 in 2D) results in numerically unstable solutions for a given lattice point because it fails to satisfy Courant-Friedrichs-Lewy condition that guarantees convergence of a forward-time central-space finite difference method 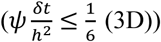. After each model run we collect the constraining variable data and in conjunction with the model parameters, we then perform RDA. Variance explained by each parameter are used to determine the influence of each parameter on the constraining variables. Refinement of the model parameters is then performed based on the variance explained and the linearity of that parameter on the constraining variables. In this way we are able to constrain the variable range iteratively by reducing the number of simulations to 5,000 (two-year simulations) while converging on parameter values. Iterations are repeated until target values are reached for each of the constraining variables of homeostasis where the combination of parameter values yields homeostasis. The secondary constraining variables served as a sanity check rather than a strict parameterization.

A pseudo-code for the parameterization process is as follows:

*Function 1: Execute Simulations given parameter ranges:*

For *n* simulations

1. Randomly select parameter values from provided ranges.
2. Run 2-year simulation of model.
3. Collect and calculate constraining variable values.

*Function 2: RDA Analysis*

While homeostasis is not reached:

1. Execute function 1 above (20,000 initially, 5,000 after).
2. Perform RDA and analyze parameter influence.
3. Further constrain ranges.
4. Repeat Step 1.

### Patient Data

Patient data was previously procured and exhaustively characterized^2^. However, the data is filtered to remove patient biopsies present in multiple biopsies as this may prove problematic in its interpretation, as previously noted^1^. The clonal area is taken as the product of the biopsy size (mm^2^) and twice the variant allele frequency. This curated dataset provides the necessary information to perform direct comparisons with simulation data.

### Model Data

Once a simulation is complete the true variant allele frequency (*VAF*_*t*_) is calculated as the quotient of the total number of cells carrying that mutation over the product of twice the total population size. This neglect copy number variation as this is not tracked within the model. However, high-throughput sequencing methodologies rarely yield the true variant allele frequency and results in imperfect information, such as that collected here. We employ and build upon a method to simulate sequence data yielding a simulated variant allele frequency (*VAF*_*s*_)^23^. The authors determine *VAF*_*s*_ by drawing the number of variant reads, *f*_*i*_ for any given mutation *i* from a binomial distribution where the success probability is given by *VAF*_*t*_ and the number of trials is itself binomially distributed and yields the total number of reads (*D*_*i*_):

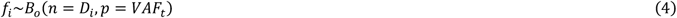

However, here *f*_*i*_ utilizes a higher resolution approach given the higher quality sequence data that is used for comparisons. Martincorena *et al.* went to great lengths to generate high resolution data by sequencing up to an average depth of 1000x (patient PD13634; while all others are approximately 500x) to ensure the capture of low frequency variants, approaching the error-rate of the sequencing method used. In order to incorporate the error and variance around the depths at each variant site that would be introduced by the sequencing methods we define *D*_*i*_ by drawing from a gamma distribution. The gamma distribution shape parameters, *K*_*p*_ and *θ*_*p*_, are specific to each patient and determined by fitting each patient’s variant depth distributions using a maximum likelihood estimation (conducted in R^21^ using the MASS package ^24^). *D*_*i*_, whose parameters are specific for each patient, for each variant becomes:

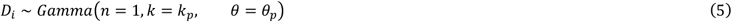

Thus, as described above, the final frequency of a variant for comparisons is given as,

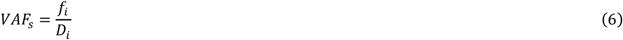

### Assessing neutrality

Most recently introduced mutations dominate the lowest frequencies and thus are difficult to observe. By using a VAF cutoff of 0.005, the lowest VAF observed in the patient data, a large fraction of mutations are filtered out and analysis is conducted on what is likely to be observable within high resolution patient sequence data.

Given a neutrally expanding keratinocyte population driven by progenitor proliferation with a basal progenitor loss/replacement rate given by *rλ*, it follows that the clone, *n*, distribution can be calculated for any given patient of age *t*:

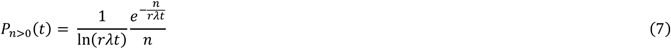

The cumulative probabilities of clone size distributions can then be used to assess neutrality, under an expectation of exponential size dependency given by the first incomplete moment^1^:

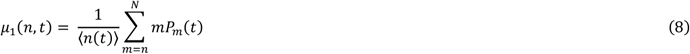

Here, *N* represents the largest clone observed and ⟨*n*(*t*)⟩ is the average mutant clone size. This assessment of neutrality allows for a straightforward method when applied to simulation data and patient biopsies. Clone size can be calculated by 2 * *VAF* * *B*, where *B* is either patient biopsy size or simulated biopsy area in *mm*^2^.

### NOTCH1 Advantage

Upon induction of a mutation within NOTCH1 a fitness advantage is given to that cell and its progeny, if any. The fitness advantage gained through a NOTCH1 mutation is non-proliferative and allows the cell to remain within the basal layer longer than it would have otherwise. This is accomplished utilizing a blocking probability, *f*_0_, which is independent for each cell regardless of clonal lineage. This indirectly impacts cell division at the basal layer, since successful division requires displacement of neighboring cells towards the corneal layer. We assessed a range of *f*_0_ values up to *f*_0_ = 1.0, where the tissue height is a quarter of its parameterized height.

### TP53 Advantages

Introducing functional heterogeneity by a TP53 advantaged clone does not involve a cell intrinsic parameter. Rather, UV damage kills cells on a given sun damage day, S, where the proportion of non-TP53 mutant cells is given by θ_s_ (Figure 2A). The TP53 mutant cells are subject to the same dynamics aside from this reprieve from death induced by UV damage. We determined the parameters to use for the sun damage day set, defined by the number of days and the spacing of those days throughout the year, and θ_s_ by approximating our 3D-HCA using an ODE that models keratinocyte population dynamics in a space limited condition.

Sun exposure varies within the lifetime of each individual. We assume that a baseline UV exposure is taken into account for the parameterization of the model, encompassing the empirically derived turnover (birth/death) of cells within the basal layer. Thus, it is necessary to have a method of UV damage and subsequent TP53 mutant advantaged clones without a direct increase in proliferation rates. The number of sun damage days within a given year defines the sun day set (S) and can take on a number of values depending on how those sun damage days are spaced throughout the year.

There are 365 possible sun damage days a year. It is unlikely that an individual would be subjected to all of these sun days, but there is a high degree of variability between individuals. In order to assess a range of values for the number of sun days within a given year, the spacing of those days, and the proportion of cells damaged by UV damage on each of those sun days we employ an ordinary differential equation (9). This approach takes a complex model, reduces its complexity, and spatial dynamics necessary to evaluate clonal dynamics in order to assess UV damage on the spatial model. We ignore the necessary information that can only be gained from a functionally heterogeneous spatial context in order to determine and implement three additional parameters to then incorporate into the spatial model (3D-HCA). The total population size over time for our homeostatic, spatial model was approximated by can be defined by an ordinary equation model that describes population growth dynamics over time in a space limited condition. The equation is

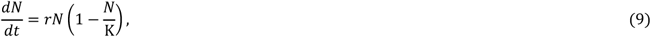

where *N* is the number of keratinocytes, *r* indicates a growth rate, *K* stands for a carrying capacity representing space limitation. Equation (9) allows us to approximate the same initializing conditions within the spatial model (3D-HCA) by initializing the model with the total number of cells in the simulated domain. Upon initialization the population reduces to the carrying capacity (*k*) over time. In order to incorporate death from UV-damage we introduce a new component:

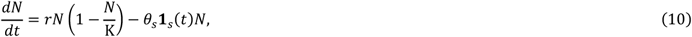

where the proportion of cells, *θ*_*s*_, at time, *t*, is killed by sun exposure. The indicator function defined as,

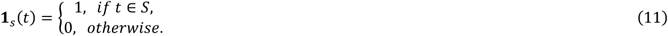

The set of sun days, *S* = {*t,t* ∈ ℤ,*t*≤ 365} because *S* ⊂ *Y* (the total days in a year), where UV damage occurs depends on the number of days chosen to repeat each year and the spacing of those days. We see that at realistic values for *θ*_*s*_ the ODE model is able to approximate our spatial simulations well, while a larger set of sun days and higher *θ*_*s*_ result in complete, or nearly complete, loss of tissue (Figure S2). In addition, the spatial model (3D-HCA) follows a logistic growth after a delayed period. This delayed period is exacerbated by higher values of *θ*_*s*_ and/or by less spacing in the number of sun days. This is the result of cells settling upon large death, creating a sparse tissue (in the absence of a TP53 mutant), the cells settle, partially, towards the basal layer prior to re-establishing homeostasis (Figure S2).

Here we assess a range of values for *θ*_*s*_ and sun day sets,*S*. The number of sun days varies for each individual as does the number of spacings between sun exposure days. We attempt to address both of these variables with regard to homeostasis. We assessed a number of sun days throughout the year (7, 20, and 100) and the spacing of those sun days (1 day to evenly spaced throughout the year). We can see that the fewer the number of sun days the least amount of tissue is lost (Figures S2 and S3). Likewise, the more spaced out these sun days are the closer to homeostasis the tissue remains (Figure S3). This is true for both the ODE and spatial models.

### Departure from Homeostasis

In order to assess how the 3D-HCA behaves in response to functional heterogeneity being introduced through NOTCH1 and TP53 mutants we use a distance metric *d*_*H*_. This distance metric summarizes the departure from homeostasis given a normal morphology such that the overall population size is constant throughout the simulation (e.g. all neutral simulations). It is defined as

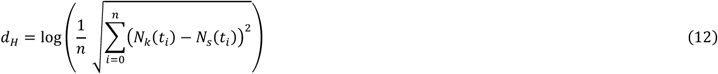

where *N*_*k*_ and *N*_*s*_ are the population size at the given time, *t*_*i*_, for the homeostatic and simulation in question, respectively. This provides a single, relative value to assess how far from homeostasis any simulation departs.

### Source Code Availability

The model was created using the HAL framework^10^ with a custom module (HAL framework available here: https://github.com/MathOnco/HAL.git). All source code specifically for this epidermis model will be made available on GitHub at the time of publication at (https://github.com/MathOnco/HomeostaticEpidermis.git).

## Supplemental Tables and Figures

**Figure S1.**
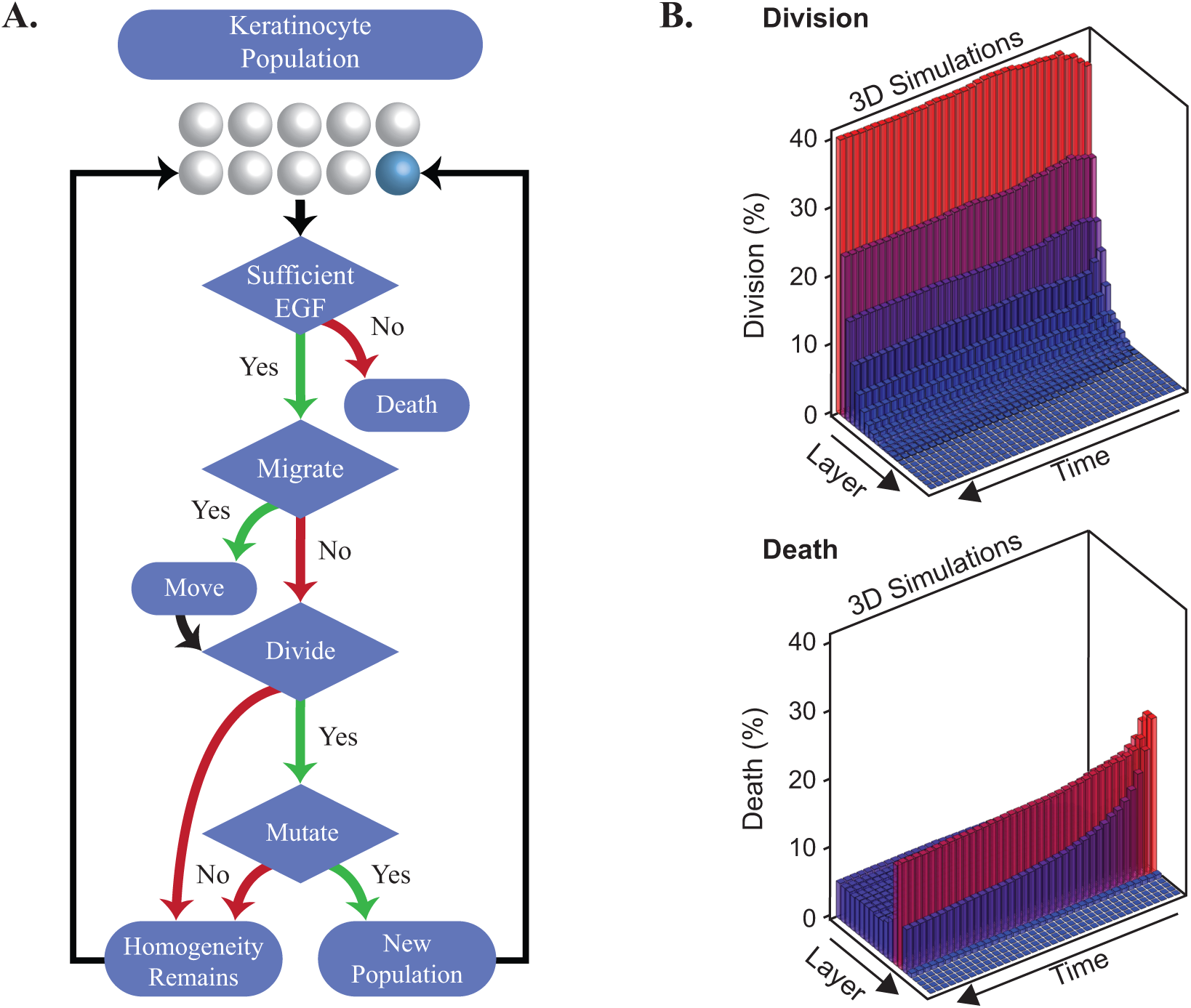
Three dimensional hybrid cellular automata keratinocyte lifecycle flowchart. (**A**) illustrates the lifecycle of a Keratinocyte cell. Prior to the keratinocyte lifecycle the EGF diffusible is solved until steady state (Equation 1 main text). Each cell step is possible during the course of the day time step. For birth and death within the model, the contributions of each are given for each layer of the epidermis over the course of a 2 year simulation (**B**).

**Table S1.** Model Parameters. Parameters are separated into the cellular automata parameters and those belonging to the diffusible, Epidermal Growth Factor (EGF).

| 3D-HCA Model Parameters |  |  |
| --- | --- | --- |
| Parameter |  | Role |
| $\alpha$ | | Level at which apoptosis can occur by chance. |
| $\theta$ | | Random probability of death. |
| $\xi$ | | Scaling factor to control proliferation rate. |
| $\varrho$ | | Probability of moving into an empty position. |
| $\omega$ | | Probability governing direction of division. |
| EGF Diffusible Parameters |  |  |
| Parameter |  | Role |
| $\psi$ | | Rate of EGF diffusion from basal layer. |
| $\lambda$ | | Rate of EGF decay. |
| $\gamma$ | | Rate of EGF consumption by keratinocytes. |

**Figure S2.**
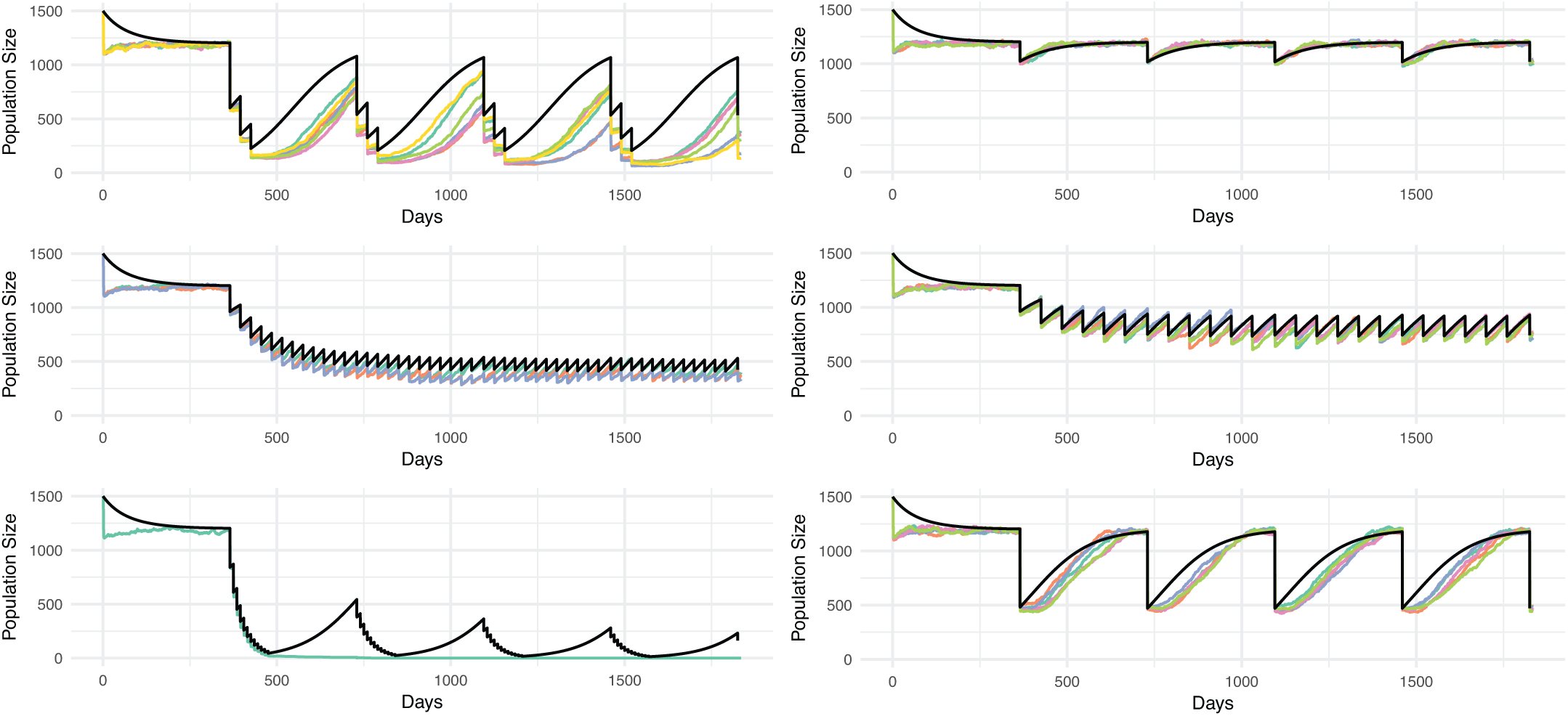
Logistic growth model is able to capture spatial dynamics to aid in evaluating TP53 sun days and dynamics. Here we define various sun day spacings, *S*, and the proportion of cells killed during sun days, *θ*_*s*_. Black line in each plot shows the ODE logistic growth model and the colored lines are for the spatial model (Methods). The first three plots on the left show severe and infrequent or, simply frequent perturbations. Whereas, for the right three plots, we see less severe perturbations through parameter combinations.

**Figure S3.**
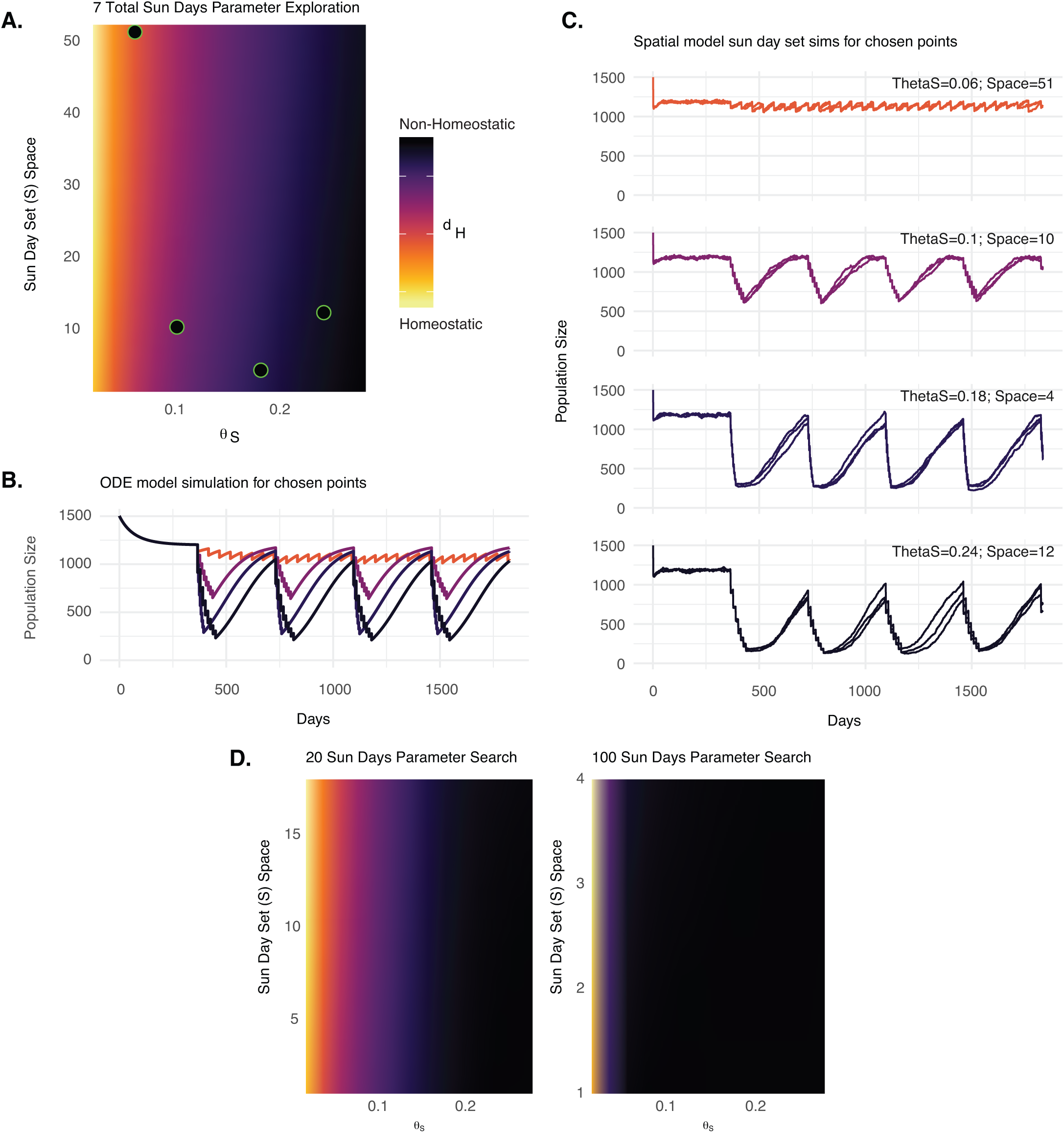
TP53 parameter evaluation using a space limited ordinary differential equation (ODE). Evaluation of the number of sun days and *θ*_*s*_ values using the ODE model. In (**A**) we see the homeostatic measure *d*_*H*_ as a response to sun day spacing and *θ*_*s*_ repeating yearly. For each black point with a green border the population sizes over time is given for the ODE (**B**) and the spatial 3D-HCA model (**C**). For (**B**) and (**C**) the colored lines are colored for where they fall within (**A**). (**D**) shows the same information, but for different numbers of sun days. The available spacing of these sun days for (**A**) and (**D**) differ due to how many days are available to space the sun days throughout the year.

## Supplemental Video Captions

**Movie S1. Neutral Model Simulation (video S1.EpidermisNeutral.mp4).** A 0.4*mm*^2^ 58 year neutral simulation. The initial and ending conditions are displayed on the upper corners of the panel. Each color represents a completely independent population (e.g. a population differing from its parent by at least one mutation). Time steps are shown and each frame represents a 6 month change.

